# Near telomere-to-telomere genome assembly of the Chilean jack mackerel *Trachurus murphyi*

**DOI:** 10.1101/2025.10.22.683973

**Authors:** Sandra Ferrada-Fuentes, Claudio Quevedo, Victoria Herrera-Yáñez, Cristian B. Canales-Aguirre, Felipe Aguilera

## Abstract

The Chilean jack mackerel (*Trachurus murphyi*) is an economically important pelagic fish widely distributed across the South Pacific Ocean. Due to intense fishing pressure and the lack of high-quality genomic resources, the development of sustainable fisheries management strategies has been hindered. Here, we present a near telomere-to-telomere genome assembly of *T. murphyi* using PacBio and Hi-C sequencing data. The assembled genome spans 818.31 Mb, with a contig N50 length of 24.89 Mb and a scaffold N50 length of 34.34 Mb. The assembly is anchored on 24 pseudo-chromosomes, covering 95.80% of the genome. BUSCO analysis indicates a high level of completeness at 99.0%, and telomeres and centromeres were identified on pseudo-chromosomes. Repetitive sequences account for 24.24% of the total genome, and 47,734 protein-coding genes were annotated, 90.98% of them with assigned function. Comparative genomics analyses reveal strong synteny with its congeneric *Trachurus* species. This high-quality genome assembly serves as a valuable resource for studying the genetic diversity and population structure of the Chilean jack mackerel and provides a solid foundation for future management and conservation efforts.

## Background & Summary

Jack mackerels are pelagic fish species of the genus *Trachurus*, found in all world’s oceans except the Arctic and Antarctic waters. They belong to the Carangidae family, a group of ray-finned fishes that includes jacks, pompanos, and trevallies^1^, and are classified into 14 species^2^. These species play a crucial role in marine ecosystems due to their trophic position in the marine food web^3^ and their importance as valuable fishery resources worldwide^4^. They are commercially exploited in many countries and are consumed fresh, frozen, smoked, or canned. Among *Trachurus* species, the Chilean jack mackerel (*T. murphyi*) fishery has been one of the largest globally, with catches peaking at nearly 5 million tons in the mid-1990s^4^. *T. murphyi* is distributed across the South Pacific Ocean^5^, allowing it to exploit a wide range of environmental conditions due to its presence in regions with highly heterogenous oceanographic and prey dynamics^6–8^. Thanks to its rich nutritional profile, *T. murphyi* is an excellent source of high-quality proteins, polyunsaturated fatty acids, vitamins, and minerals^9^. It could be that for those reasons, this species has been harvested for human consumption since at least the Early Holocene^10,11^.

Over the past decades, the Chilean jack mackerel population has declined significantly due to a combination of factors, including overfishing and variability with the El Niño-Southern Oscillation (ENSO)^4,12–15^. Consequently, the implementation of effective fisheries management strategies is crucial. Previous studies have explored the specieś fishery biology and genetics^16–20^. Population genetic analyses have revealed that *T. murphyi* constitutes a single, large panmictic population across its entire geographic range^18,21–24^. Additionally, recent research has identified a ZW sex determination system in *T. murphyi*^25^. Despite these findings, genomic and transcriptomic resources for *T. murphyi* remain scarce, with the mitochondrial genome being only recently described resource^26^. More broadly, genomic data for jack mackerels (*Trachurus*) are limited, with genome sequences currently available only for the Atlantic jack mackerel (*Trachurus trachurus*)^27^ and the Japanese jack mackerel (*Trachurus japonicus*)^28^.

In this study, we combined PacBio long reads and high-resolution chromosome conformation capture (Hi-C) sequencing to generate a near telomere-to-telomere genome assembly of *T. murphyi*. The final assembly spans 818.31 Mb, with a contig N50 of 24.89 Mb and a scaffold N50 of 34.34 Mb, comprising 24 pseudo-chromosomes and achieving 95.80% assembly coverage. The completeness assessment using Benchmarking Universal Single-Copy Orthologs (BUSCO) indicates that 98.8% of conserved metazoan and 99.0% actinopterygian genes are present and complete. The *T. murphyi* genome encodes 47,734 protein-coding genes, with 90.98% (43,427 genes) annotated with functional information. Comparative genomic analysis among *T. murphyi, T. trachurus*, and *T. japonicus* genomes reveals a high degree of synteny, suggesting limited divergence since their last common ancestor, which existed approximately ±20 mya^2^. This near telomere-to-telomere genome assembly provides a valuable resource for advancing our understanding of the genomic structure and genetic traits of *T. murphyi*, laying a solid foundation for future fishery management and conservation strategies. Additionally, this study offers new insights into the biological and ecological adaptation mechanisms of jack mackerels and underscores the importance of exploring genomic and ecological differences among closely related species, extending the relevance *Trachurus* genomics beyond fisheries management.

## Methods

### Sample preparation and DNA sequencing

An adult Chilean jack mackerel (fork length of 45.4 cm and weights 1.395 kg; Fig. 1A) was obtained from commercial fisheries in the Biobío Region, Chile (36.94°S, 73.75°E) in June 2022. Muscle tissue was dissected and immediately stored at -80°C until shipped on dry ice to Cantata Bio (https://cantatabio.com/) for DNA extraction, genomic DNA sequencing, and Hi-C library construction.

**Figure 1.**
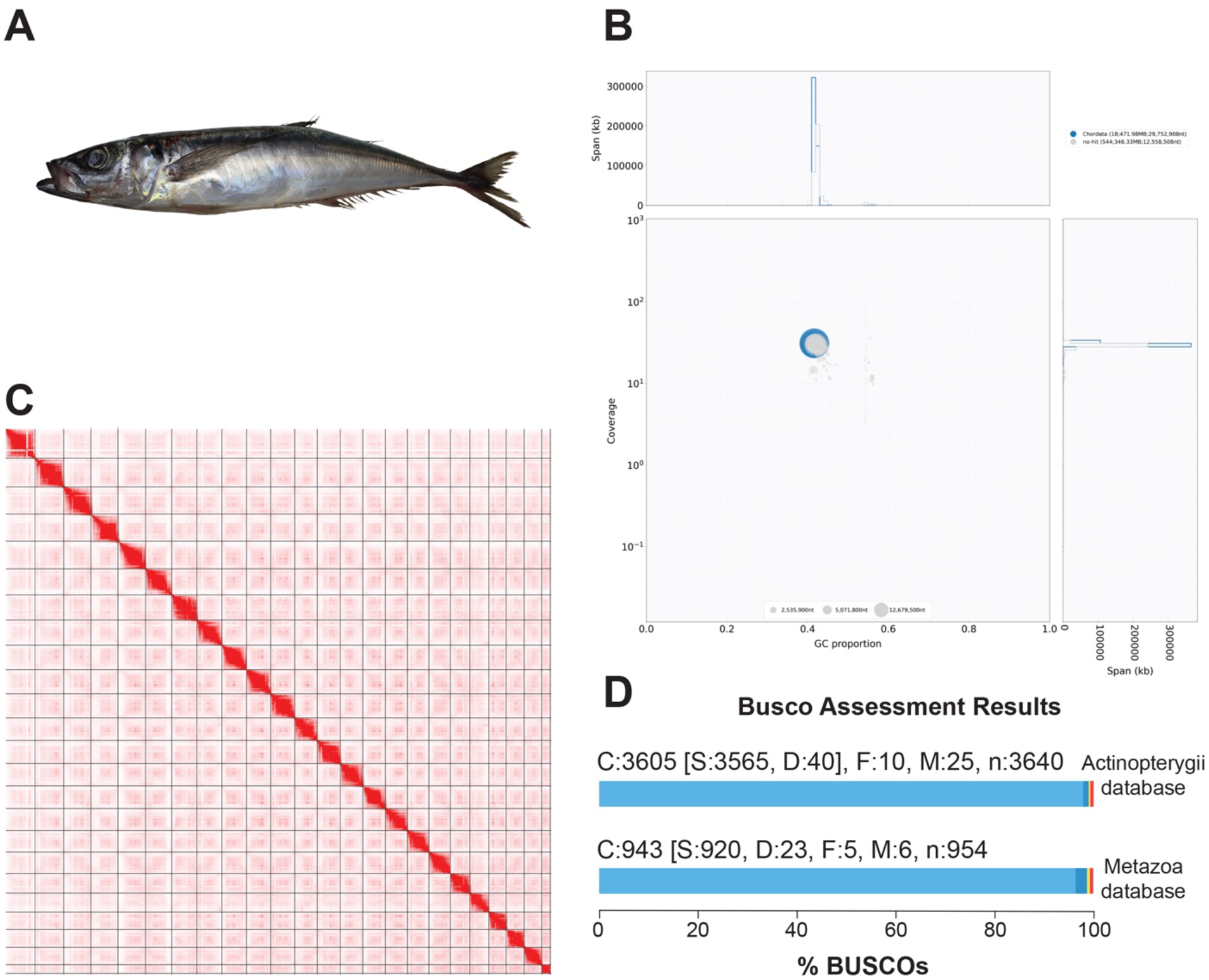
Chromosome-level genome assembly of *Trachurus murphyi*. (**A**) A photograph of the *T. murphyi* used for whole genome sequencing. (**B**) Square-binned blob plot showing the distribution of assembly contigs on GC proportion and coverage axes. Squares within each bin are colored according to taxonomic annotation and scaled accordingly to total span. (**C**) The contact map of Omni-C interactions for the assembly after genome scaffolding. (**D**) Histogram of BUSCO assessment of the *T. murphyi* genome against Actinopterygii and Metazoa databases.

High-quality DNA was used to construct two PacBio Hi-Fi SMRTbell^®^ libraries and one Dovetail Omni-C^®^ sequencing library for assembling the *T. murphyi* genome. The PacBio libraries were sequenced with circular consensus sequencing (CCS) mode on a PacBio Sequel II system, generating 7,058,275 High-Fidelity (HiFi) reads, totaling 86.5 Gb of data, which covered the genome at 108-fold, with a N50 read length of 12,194 bp. For the Dovetail Omni-C^®^ library, chromatin was first crosslinked with formaldehyde within the nucleus before extraction. The fixed chromatin was then digested with DNAseI, followed by end repair and ligation to a biotinylated bridge adapter, enabling proximity ligation of adapter-containing ends. After ligation, crosslinks were reversed, and the DNA was purified. Biotin that was not internal to ligated fragments were removed, and the sequencing library was prepared using NEBNext Ultra enzymes and Illumina-compatible adapters. Biotin-containing fragments were enriched using streptavidin beads before PCR amplification. The final library was sequenced on an Illumina HiSeqX platform using 2 x 150 bp reads, generating 87,721,376 read pairs, amounting to 28.6 Gb of data, with a coverage of 32.1-fold across the genome.

### Chromosome-level genome assembly with PacBio and Hi-C data

The PacBio HiFi reads were initially assembled using Hifisiam^29^ (v0.15.4-r347) with default parameters to generate both the primary and haplotigs-alternative sequences for regions where heterozygosity was detectable within primary contigs using long reads (Table 1). The assembled contigs were then screened for potential contamination using the BlobToolKit^30^ (v4.4.3) and the BLAST non-redundant database (Fig. 1B). Contigs identified as potential contamination were removed from the assembly (Table 1). Finally, purge_dups^31^ (v1.2.5) was applied to eliminate haplotigs and overlapping contigs.

**Table 1.** Summary statistics of the assembled contigs and scaffolds of *Trachurus murphyi*.

| Features | Hifiasm<br>assembly | Hifiasm +<br>BlopToolKit<br>assembly | Purge_dups<br>assembly | Dovetail HiRise<br>assembly |
| --- | --- | --- | --- | --- |
| Total sequences | 562 | 562 | 337 | 523 |
| Total bases (bp) | 818,306,126 | 818,306,126 | 802,744,728 | 818,265,206 |
| Longest sequence (bp) | 50,717,993 | 50,717,993 | 50,717,993 | 43,616,551 |
| N50 length (bp) | 24,885,125 | 24,885,125 | 24,885,125 | 34,335,234 |
| N90 length (bp) | 6,328,993 | 6,328,993 | 6,633,185 | 25,125,839 |
| Ns (%) | 0.00 | 0.00 | 0.00 | 0.00 |

Following these steps, the contig-level genome assembly spans 802.27 Mb, comprising 337 contigs with an N50 length of 24.89 Mb (Table 1).

To achieve a chromosome-level assembly, we used the contig-level genome assembly and Dovetail OmniC^®^ reads as input for HiRise, a software pipeline specifically designed to scaffold genome assemblies using proximity ligation data^32^. Briefly, Dovetail OmniC® reads were mapped to the contig-level genome assembly using BWA (https://github.com/lh3/bwa) with a minimum quality threshold of 50 (MQ>50). HiRise then analyzed the separation of Dovetail OmniC^®^ read pairs mapped within draft scaffolds to generate a likelihood model for genomic distances. This model was used to identify and correct putative misjoins, score prospective joins, and execute those joins above a set threshold. Following these steps, we generated a Hi-C contact map and manually reviewed it using Juicebox^33^ (v1.11.08) to correct any misjoins and position unplaced contigs. This process resulted in a high-quality, chromosome-level genome assembly with a total length of 818.31 Mb and an N50 length of 34.34 Mb (Table 1). The Hi-C analysis upgraded the contig-level assembly to a chromosome-level assembly comprising 24 chromosomal sequences (Fig. 1C). These 24 sequences covered 95.80% of the genome, while the remaining unplaced scaffolds accounted for 34.31 Mb (Table 2).

**Table 2.** Chromosome lengths and telomere statistics of *Trachurus murphyi*.

| Chromosome name | Length (bp) | GC (%) | Telomere | Left length (bp) / Right length (bp) of telomere sequence |
| --- | --- | --- | --- | --- |
| Chr1 | 43,616,551 | 42.17 | No | 0 / 0 |
| Chr2 | 40,697,839 | 41.62 | No | 0 / 0 |
| Chr3 | 39,753,857 | 41.81 | Left | 3,420 / 0 |
| Chr4 | 39,403,342 | 41.91 | Right | 0 / 4,326 |
| Chr5 | 38,686,146 | 42.20 | No | 0 / 0 |
| Chr6 | 36,848,658 | 42.03 | Right | 0 / 1,782 |
| Chr7 | 36,663,105 | 41.79 | No | 0 / 0 |
| Chr8 | 36,082,001 | 42.16 | Right | 0 / 1,362 |
| Chr9 | 35,151,152 | 42.16 | No | 0 / 0 |
| Chr10 | 34,807,739 | 42.01 | Left | 1,212 / 0 |
| Chr11 | 34,335,234 | 41.94 | No | 0 / 0 |
| Chr12 | 33,135,089 | 42.20 | Left | 5,460 / 0 |
| Chr13 | 32,997,167 | 41.80 | No | 0 / 0 |
| Chr14 | 32,784,410 | 41.70 | No | 0 / 0 |
| Chr15 | 32,188,078 | 42.45 | No | 0 / 0 |
| Chr16 | 30,992,529 | 42.29 | Right | 0 / 3,660 |
| Chr17 | 30,849,155 | 41.81 | No | 0 / 0 |
| Chr18 | 30,703,906 | 41.91 | Left | 1,266 / 0 |
| Chr19 | 28,653,432 | 42.02 | Left | 828 / 0 |
| Chr20 | 27,799,838 | 42.19 | Right | 0 / 1,104 |
| Chr21 | 25,177,033 | 42.28 | Right | 0 / 5,094 |
| Chr22 | 25,125,839 | 42.19 | Right | 0 / 5,760 |
| Chr23 | 24,528,291 | 42.13 | No | 0 / 0 |
| Chr24 | 12,959,388 | 43.22 | No | 0 / 0 |
| Unplaced scaffolds | 34,308,866 | 48.44 | - | - |
| ChrMT | 16,561 | 46.59 | - | - |

Finally, the completeness of the chromosome-level genome was assessed using BUSCO^34^ (v5.3.2), revealing that 99.0% and 98.8% of actinopterygian and metazoan conserved genes are present in the final assembly (Fig. 1D).

### Identification of telomere and centromere regions in the chromosome-level genome assembly

Given the high quality and gapless nature of our chromosome-level genome assembly (Table 1), we sought to determine whether it meets the criteria for a near telomere-to-telomere genome assembly. To identify telomeric regions, we searched the Chilean jack mackerel genome for the (TTAGGG)n sequence, the canonical telomere repeat found in most animals^35^. Telomere identification was carried out using the TeloExplorer function of the quarTeT^36^ (v1.2.5) software with the “-c animal” option. Our analysis revealed telomere regions with high confidence at one end of 12 chromosomes (Table 2). A visual representation of the telomere distribution is provided in Fig. 2A.

**Figure 2.**
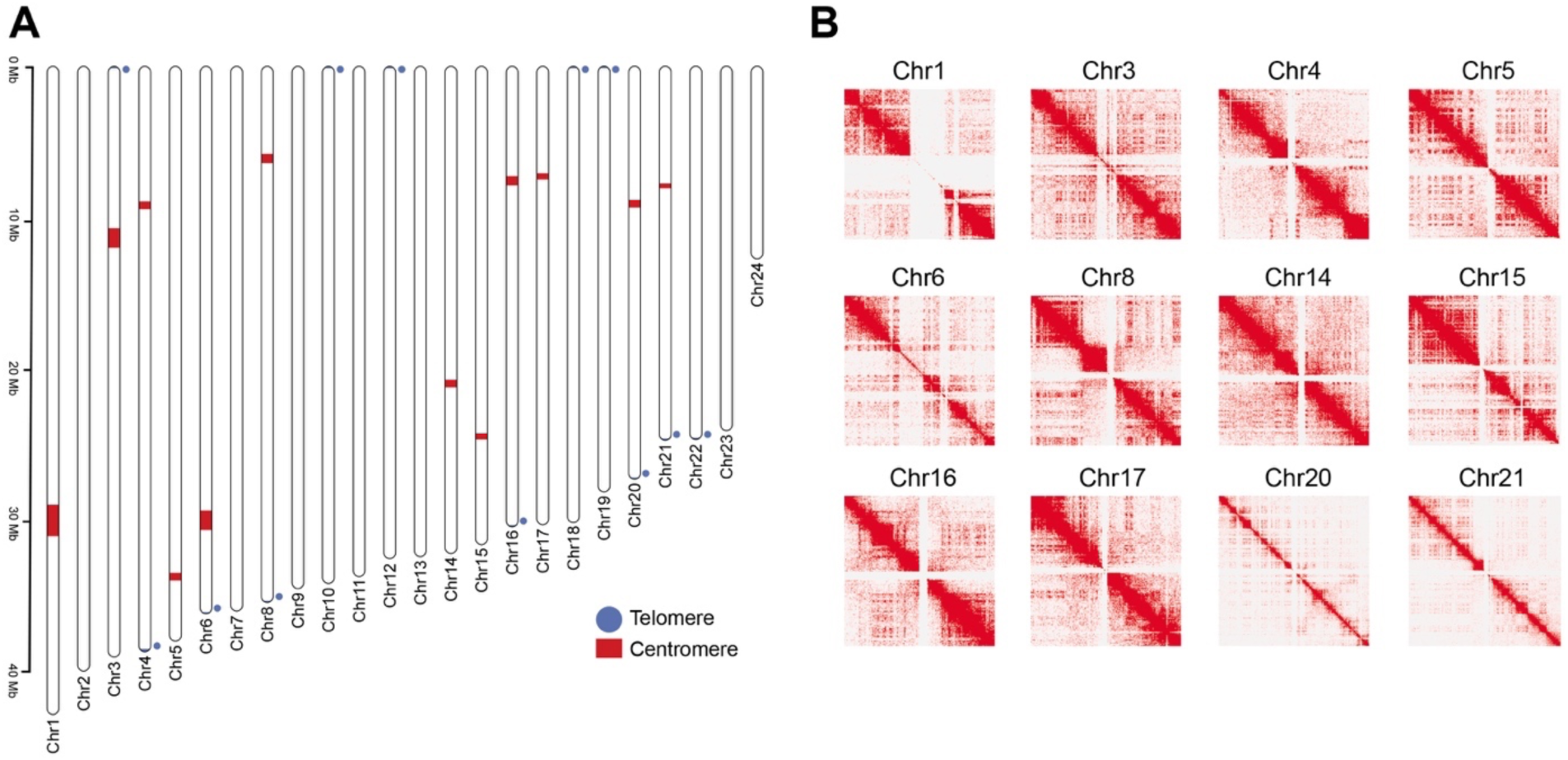
An overview of the telomeres and centromeres of the near telomere-to-telomere genome assembly of *Trachurus murphyi*. (**a**) Chromosome ideograms showing the telomeric (circles) and centromeric (rectangles) regions identified by TeloExplorer and CentroMiner modules in quarTeT^36^, respectively. Telomeric regions are colored in blue, while centromeric regions are colored in red. (**b**) Omni-C chromatin interactions at 50 kb resolution reveals the characteristics of the centromere region in the Chilean jack mackerel genome.

To identify centromeric regions, which are characterized by large stretches of repetitive satellite DNA or tandem repeats ranging in size from hundreds of kilobases to megabases^37^, we used the CentroMiner tool of the quarTeT^36^ (v1.2.5) software with default parameters. Briefly, this tool detects all tandem repeat monomers in a genome assembly and selects those most likely to be centromeric repeats on each chromosome based on their period and copy number. These monomers are then clustered to reduce redundancy and aligned to the corresponding chromosomes. Continuous matching regions are added to the candidates set, and scores are assigned based on total match length and retrotransposon content to ultimately determine the locations and predominant monomer repeats^36^. Our findings revealed that putative centromeric regions range in length from 0.3 Mb to 2.1 Mb (Fig. 2A, Table 3). Notably, these regions contain few protein-coding genes but are rich in dispersed repeats, with LTR/Gypsy, LINE/L2 and simple repeats being the predominant types (Table 3), which is in concordance with typical centromere definition^38^. We also observed large blank regions in the Omni-C interaction heatmap of the centromere region (Fig. 2B), a characteristic feature of centromeres that have also been observed in other gap-free genomes where centromeres have been identified^39,40^. The locations of the centromeres are indicated on chromosome ideograms, as shown in Fig. 2A.

**Table 3.** Summary statistics and presence of repetitive sequences in the putative centromere regions of *T. murphyi*.

| Chromosome | Start position (bp) | End position (bp) | Length (Mb) | LTR/Gypsy (%) | LINE/L2 (%) | Simple repeats (%) | # of genes |
| --- | --- | --- | --- | --- | --- | --- | --- |
| Chr1 | 29,500,000 | 31,600,000 | 2.1 | 1.73 | 1.24 | 0.0 | 112 |
| Chr2 | - | - | - | - | - | - | - |
| Chr3 | 10,900,000 | 12,200,000 | 1.3 | 2.57 | 6.93 | 0.0 | 63 |
| Chr4 | 9,100,000 | 9,600,000 | 0.5 | 4.22 | 0.03 | 0.11 | 16 |
| Chr5 | 34,100,000 | 34,600,000 | 0.5 | 1.79 | 0.65 | 0.43 | 19 |
| Chr6 | 29,900,000 | 31,200,000 | 1.3 | 3.30 | 1.30 | 0.0 | 104 |
| Chr7 | - | - | - | - | - | - | - |
| Chr8 | 5,900,000 | 6,500,000 | 0.6 | 3.81 | 0.51 | 0.38 | 26 |
| Chr9 | - | - | - | - | - | - | - |
| Chr10 | - | - | - | - | - | - | - |
| Chr11 | - | - | - | - | - | - | - |
| Chr12 | - | - | - | - | - | - | - |
| Chr13 | - | - | - | - | - | - | - |
| Chr14 | 21,100,000 | 21,600,000 | 0.5 | 1.14 | 0.27 | 0.0 | 9 |
| Chr15 | 24,700,000 | 25,100,000 | 0.4 | 0.92 | 0.19 | 0.68 | 12 |
| Chr16 | 7,400,000 | 8,000,000 | 0.6 | 0.62 | 0.22 | 1.24 | 26 |
| Chr17 | 7,200,000 | 7,600,000 | 0.4 | 0.97 | 0.24 | 0.81 | 23 |
| Chr18 | - | - | - | - | - | - | - |
| Chr19 | - | - | - | - | - | - | - |
| Chr20 | 9,000,000 | 9,500,000 | 0.5 | 1.65 | 0.30 | 0.22 | 28 |
| Chr21 | 7,900,000 | 8,200,000 | 0.3 | 1.76 | 0.03 | 1.22 | 8 |
| Chr22 | - | - | - | - | - | - | - |
| Chr23 | - | - | - | - | - | - | - |
| Chr24 | - | - | - | - | - | - | - |

Taken together, these results suggest that the Chilean jack mackerel genome qualifies as a near telomere-to-telomere assembly category.

### Repetitive sequence annotation

To identify repetitive sequences in the Chilean jack mackerel genome, we generated a custom repeat library using a combination of *de novo* and homology-based approaches. This included RepeatModeler^41^ (v2.0.1), TransposonPSI^42^ (v1.0.0), LTRharvest^43^ (v1.6.2), LTRdigest^44^ (v1.6.2), TIRvish^45^ (v1.6.2), MITE Tracker^46^ (v1.0), and HelitronScanner^47^ (v1.1), all executed with default parameters. The individual repeat libraries were concatenated, and sequences sharing more than 80% similarity were merged to remove redundancy using USEARCH^48^ (v11.0.667). The resulting non-redundant repeat library was then classified with RepeatClassifier^41^ (v2.0.1). To filter out protein-coding sequences, repetitive elements were mapped against a custom database, containing the annotated protein-coding sequences of *Danio rerio*, *Seriola lalandi dorsalis*, *Cyclopterus lumpus*, *Xiphophorus couchianus*, *Oncorhynchus tshawytscha*, and *Salmo salar*, using BLASTn^49^ (v2.9.0+) with an e-value threshold of 0.001. Any sequences matching protein-coding genes were removed from the library. The final repeat library was then used in RepeatMasker^50^ (v4.1.1) to generate a genome report repeat content and to mask the genome for subsequent annotation with the BRAKER3 pipeline.

We identified a total of 198.35 Mb of repetitive sequences, representing 24.24% of the assembled Chilean jack mackerel genome (Table 4). DNA transposons and retroelements are the predominant transposable elements, accounting for 8.89% and 8.34% of the genome, respectively (Fig. 3A, Table 4). Among these, the most abundant types by copy number are the hAT-Ac DNA transposon and the L2 (LINE), Gypsy (LTR), and tRNA (SINE) retroelements (Fig. 3B). Most transposable element activity appears to be relatively recent, as a large proportion of repeats exhibit a low genetic distance from their respective consensus sequences (Fig. 3C). Additionally, there is strong evidence of very recent activity in LINEs and LTRs, with a sharp increase in the number of elements exhibiting minimal genetic distance (Fig. 3C). Finally, the distribution of transposable elements suggests that DNA transposons and LTRs are enriched at the ends of the chromosomes, whereas LINEs are more evenly distributed across the chromosomes (Fig. 3D). SINEs and Rolling-circles (Helitrons) are restricted to particular chromosomes with a punctuated distribution (Fig. 3D).

**Figure 3.**
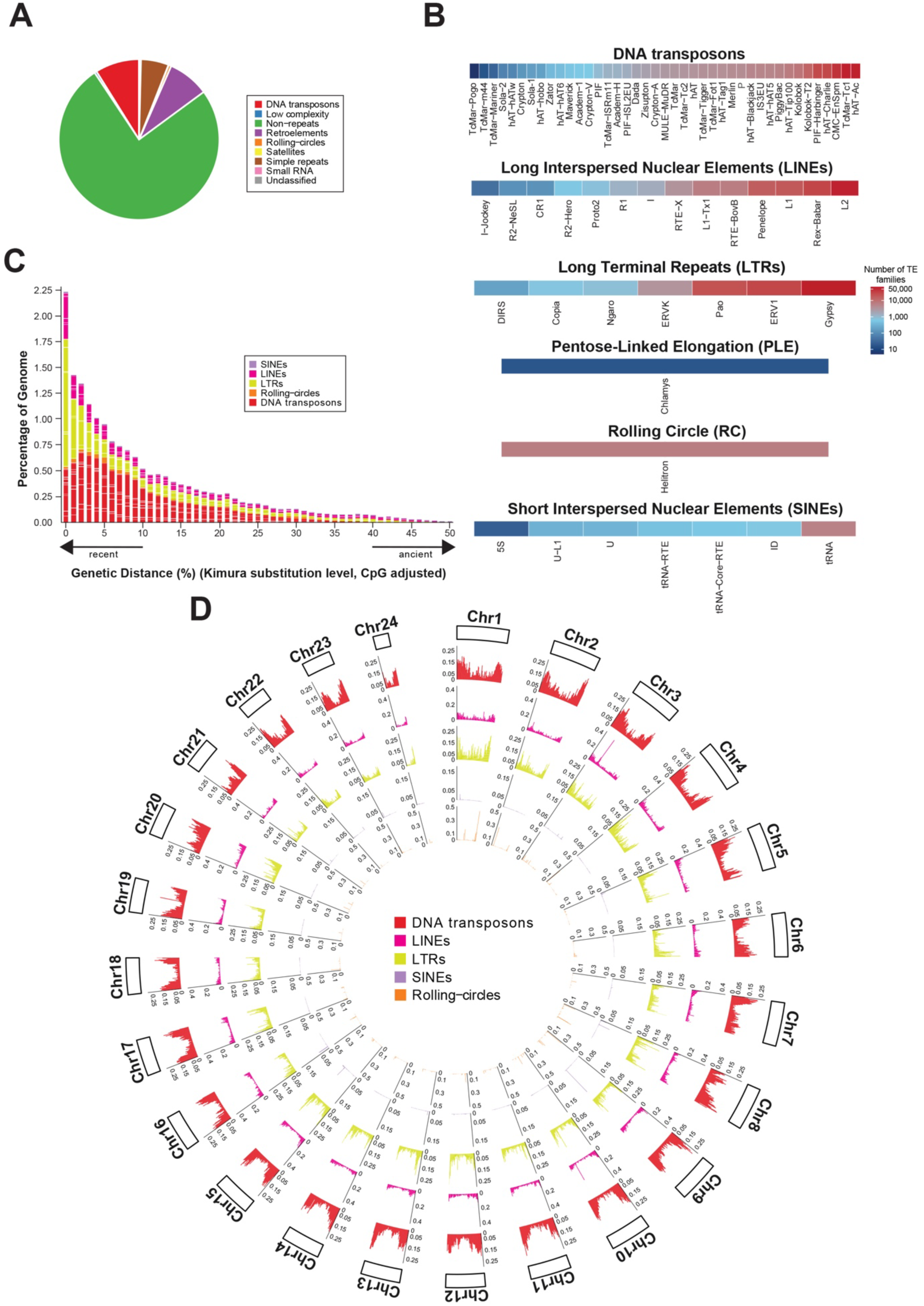
Transposable elements (TE) within the genome of *Trachurus murphyi*. (**a**) The proportion of the assembly comprised of the main TE classifications, as represented by the colors in the key. (**b**) Hierarchical clustering of the number of TE species within families. (**c**) Repeat landscape plot illustrating the proportion of repeats in the genome, with the y-axis representing the genomic coverage and the x-axis cpG-corrected Kimura divergence to the repeat consensus. (**d**) Circos plot depicting the distribution of DNA transposons, LINEs, LTRs, SINEs, and Roling-circles (Helitrons) across the chromosomes of *T. murphyi*.

**Table 4.** Repeat content and their proportion in the *T. murphyi* genome.

| Type of repetitive element | Length (bp) | Percentage (%) |
| --- | --- | --- |
| Retroelements | 68,263,474 | 8.34 |
| SINEs | 1,035,585 | 0.13 |
| L2/CR1/Rex | 16,015,217 | 1.96 |
| R1/LOA/Jockey | 686,884 | 0.08 |
| R2/R4/NeSL | 362,068 | 0.04 |
| RTE/Bov-B | 2,135,235 | 0.26 |
| L1/CIN4 | 4,627,191 | 0.57 |
| LTR elements | 39,300,853 | 4.80 |
| BEL/Pao | 7,663,374 | 0.94 |
| Ty1/Copia | 269,409 | 0.03 |
| Gypsy/DIRS1 | 21,394,851 | 2.61 |
| Retroviral | 8,882,452 | 1.09 |
| DNA transposons | 72,717,968 | 8.89 |
| hobo-Activator | 20,174,923 | 2.47 |
| Tc1-IS630-Pogo | 17,547,274 | 2.14 |
| MULE-MuDR | 681,946 | 0.08 |
| PiggyBac | 806,279 | 0.10 |
| Tourist/Harbinger | 4,772,178 | 0.58 |
| Other (Mirage, P-element, Transib) | 968,348 | 0.12 |
| Rolling-circles (Helitrons) | 4,329,195 | 0.53 |
| Unclassified | 100,161 | 0.01 |
| Total interspersed repeats | 141,085,620 | 17.24 |
| Small RNA | 3,105,178 | 0.38 |
| Satellites | 508,595 | 0.06 |
| Simple repeats | 45,640,029 | 5.58 |
| Low complexity | 3,683,752 | 0.45 |

### Gene prediction and functional annotation

To annotate protein-coding sequences in the Chilean jack mackerel genome, we used the BRAKER3^51^ (v3.0.8) along with reference proteins from 18 species: *Anabas testudineus* (GCA_900324465.3), *Carassius auratus* (GCA_003368295.1), *Clupea harengus* (GCA_900700415.2), *Danio rerio* (GCA_000002035.4), *Gadus morhua* (GCA_902167405.1), *Gasterosteus aculeatus* (GCA_016920845.1), *Oreochromis aureus* (GCA_013358895.1), *Oreochromis niloticus* (GCA_001858045.3), *Oryzias javanicus* (GCA_003999625.1), *Poecilia formosa* (GCA_000485575.1), *Salmo salar* (GCA_905237065.2), *Sander lucioperca* (GCA_008315115.1), *Scleropages formosus* (GCA_900964775.1), *Seriola dumerili* (GCA_002260705.1), *Seriola lalandi dorsalis* (GCA_002814215.1), *Sparus aurata* (GCA_900880675.1), *Trachurus trachurus* (GCA_905171665.1), and *Xiphophorus couchianus* (GCA_001444195.1). All protein sequences were obtained from Ensembl Release 112.

Briefly, GeneMark-EPT^52^ was used to identify genomic loci where extrinsic evidence support reliable gene predictions. These high-confidence predictions were then used to train the statistical model implemented in AUGUSTUS^53^. Further refinements incorporated protein-derived extrinsic evidences from the reference proteomes listed above, with gene predictions subsequently optimized using TSEBRA^54^. As a result, BRAKER3 identified 47,734 protein-coding genes, with an average gene length of 7.01 kb and an average coding sequence (CDS) length of 1.53 kb (Fig. 4A). On average, each gene contained 4.94 exons (Fig. 4B), while exon length averaging 898 bp and intron lengths averaging 977 bp (Fig. 4C). Functional annotation of the predicted protein-coding genes was conducted using DIAMOND^55^ (v2.1.11) against the following public databases: NCBI non-redundant (nr), SwissProt, TrEMBL, and eggNOG. Protein domain searches were performed using Pfam with HMMER^56^ v(3.3.2) and InterProScan^57^ (v5.74.105). Gene Ontology (GO) terms were extracted from the InterProScan output, and KEGG Orthology (KO) annotation was conducted using KofamScan^58^ and the KOfam database. Overall, 43,427 (90.98%) protein-coding genes had at least one homologous hit across these public databases, with 9,080 (19.02%) showing matches in at least eight databases (Fig. 4D).

**Figure 4.**
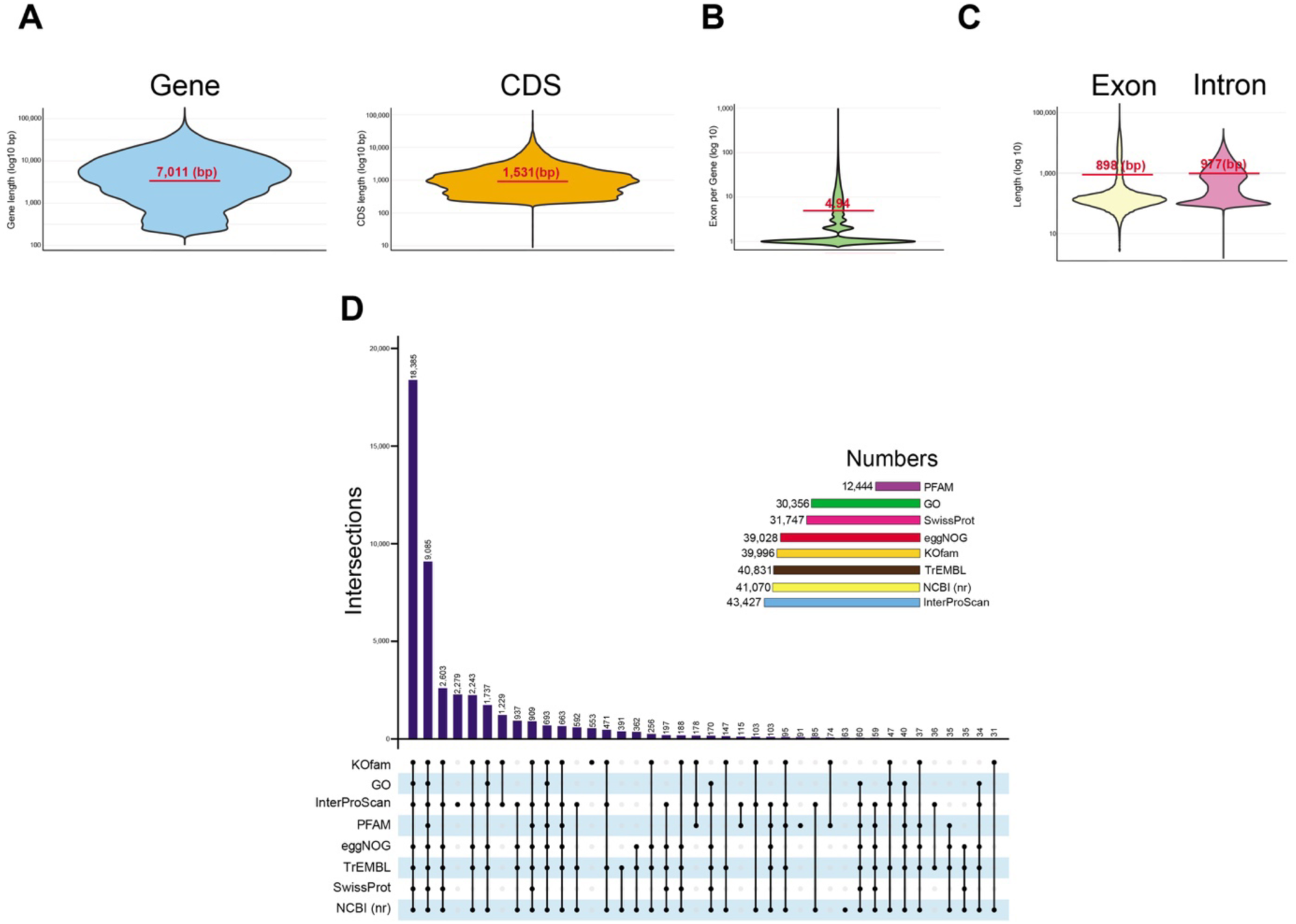
Characteristics and functional annotation of the predicted protein-coding genes in the *Trachurus murphyi* genome. (**a**) Distribution of gene lengths and coding sequence (CDS) lengths for the predicted protein-coding genes. (**b**) Violin plot showing the number of exons per gene. (**c**) Distribution of exon and intron lengths across the protein-coding genes. (**d**) UpSet diagram summarizing the functional annotation of the protein-coding genes based on eight public databases: NCBI nr, SwissProt, TrEMBL, eggNOG, Pfam, InterProScan, GO, and KOfam.

### Mitogenome assembly and annotation

To assemble the mitochondrial (mtDNA) genome of *T. murphyi*, we employed a strategy that utilizes the reference mitogenome of a closely related species from NCBI (accession number PP935111.1 in this case). This reference was used to capture mitochondrial PacBio HiFi reads, which were then assembled into a single circularized contig using MitoHiFi^59^ (v3.2.1). In addition to the annotation provided by MitoHiFi, the assembled mitogenome was further annotated using MITOS2 web server^60^. To evaluate the depth of coverage, all PacBio HiFi reads were mapped back to the assembled mitogenome using minimap2^61^ v(2.29). We used BLAST+^49^ (v2.10.1) to identify matches between the mitochondrial genome assembly and the nuclear genome assembly. Nuclear contigs showing >99% sequence identity and shorter lengths than the mitochondrial genome assembly were filtered out. The resulting mitogenome assembly and its annotations were visualized using the Proksee web tool^62^. Mitochondrial protein-coding gene arrangements among *Trachurus* species were compared using MITOS2 GFF annotation data.

The assembled mitogenome of *T. murphyi* is 16,561 bp in length and comprises 13 protein-coding genes, two rRNA genes, 22 tRNA genes, and successfully circularized (Fig. 5A). Coverage analysis revealed relatively uniform read depth across most of the mitochondrial genome, except for two notable regions: reduced coverage in the first 5 kb, likely associated with repetitive or low-complexity elements, and a pronounced drop in coverage near the end of the linearized genome representation (Fig. 5B). This terminal decrease is a common artifact of mapping reads to a linearized circular genome, as reads spanning the origin or control region may not align properly.

**Figure 5.**
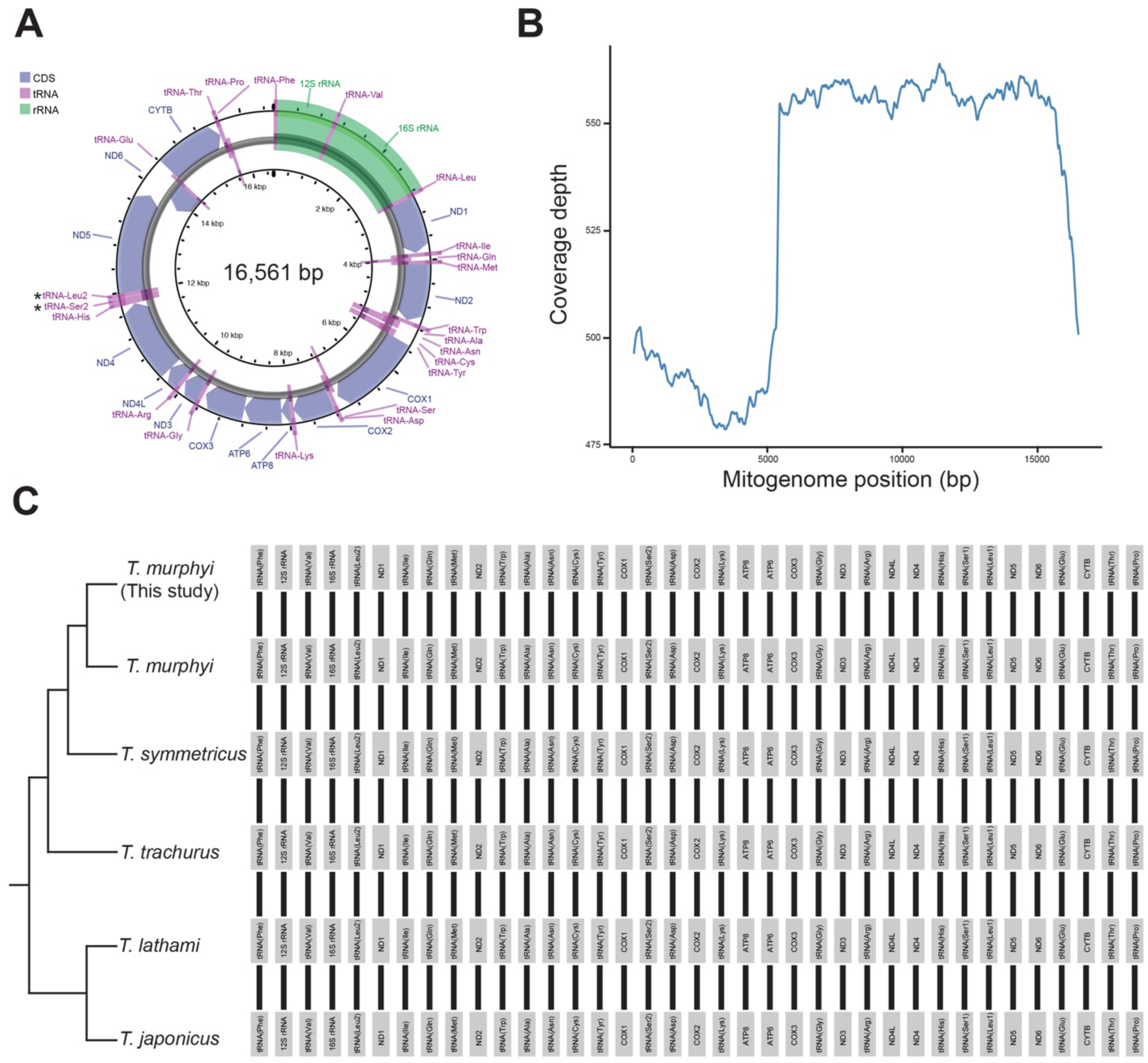
Mitochondrial genome of *Trachurus murphyi* and comparison with other *Trachurus* species. (**a**) Organization of the *T. murphyi* mitochondrial genome visualized as a Circos plot. (**b**) Coverage depth across the mitogenome based on PacBio HiFi reads. (**c**) Collinearity analysis of six mitogenomes from *Trachurus* species. Homologous regions between mitogenomes are connected by black lines. Dendrogram to the right depicts phylogenetic relationships among *Trachurus* species.

Collinearity analysis revealed no variation in the mitochondrial genomes among six *Trachurus* species (Fig. 5C), suggesting strong evolutionary constraints on gene arrangement due to the close relatedness of the species analyzed. This finding is consistent with a recent common ancestor that diverged approximately 20 mya^2^.

### Comparative genomics between *Trachurus* species

To assess the extent of genomic conservation among *Trachurus* species, we compared genome assemblies among *T. murphyi* (this study), *T. trachurus*^27^ (NCBI assembly accession: GCA_905171665.2), and *T. japonicus*^28^ (NCBI assembly accession: GCA_045865225.1). Long sequences (>10 Mbp) corresponding to pseudo-chromosomes were extracted from each genome assemblies to facilitate clear chromosome-level comparisons. To identify potential structural rearrangements, we used D-GENIES^63^ to align and compare the chromosomes of the three *Trachurus* species. The alignments revealed a high degree of synteny, despite several structural rearrangements. In *T. trachurus*, we observed inversions of 0.6-4.7 Mbp and duplications of 4.1-5.1 Mbp (Fig. 6A). Similarly, *T. japonicus* exhibited inversions (0.5-3.1 Mbp), duplications (1.1-5.6 Mbp), and translocations (1.7-3.4 Mbp) (Fig. 6B).

**Figure 6.**
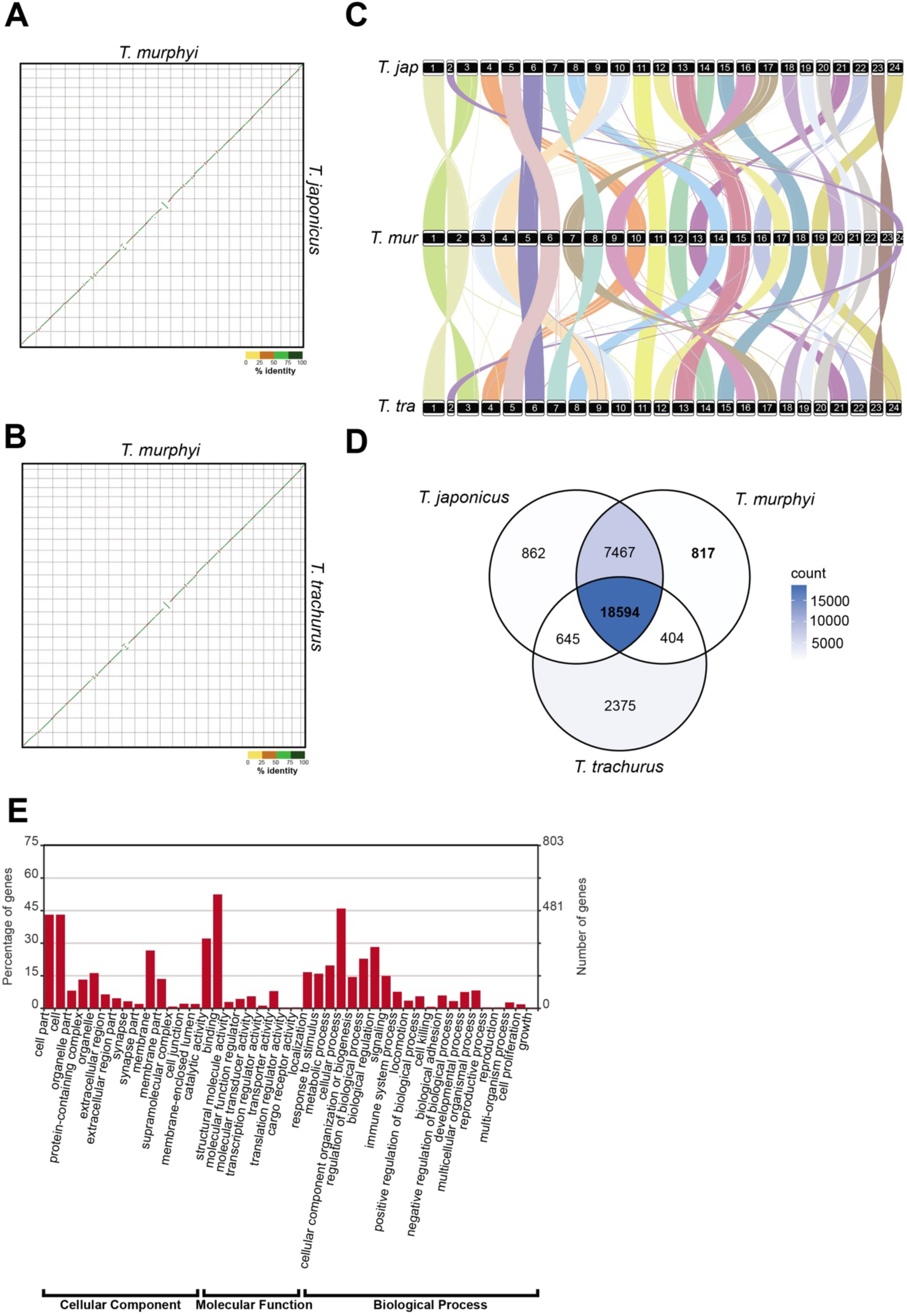
Comparative genomics among *Trachurus* species. (**a**) Oxford dot plot showing the genomic coordinates between *T. japonicus* and *T. murphyi*. Chromosome comparisons are color-coded based on the percentage of sequence identity. (**b**) Oxford dot plot showing the genomic coordinates between *T. murphyi* and *T. trachurus*. Chromosome comparisons are color-coded based on the percentage of sequence identity. (**c**) Synteny (gene order) comparison among *Trachurus* species. Orthologous genes with a 1:1:1 relationship across the three species are shown in different colors. Color blocks represent genes located on specific chromosomes, indicated by numbers within the squares. Tjap = *Trachurus japonicus*; Tmur = *Trachurus murphyi*; and Tra = *Trachurus trachurus*. (**d**) Venn diagram illustrating shared and unique gene families among the three *Trachurus* species. The total number of gene families for each species is indicated. (**e**) GO distribution of gene families unique to *T. murphyi*, categorized by cellular compartments, molecular functions, and biological processes

Nevertheless, our results suggest strong evolutionary conservation at the genome-wide level among *Trachurus* species.

Orthologous gene clusters in a 1:1:1 relationship were identified using Orthofinder^64^ (v2.3.8) with an e-value cut-off 1e^-5^, employing an all-to-all DIAMOND analysis. Comparing the genomic locations of 10,844 orthologous genes across the three genomes revealed disrupted gene clusters (Fig. 6C). Despite these rearrangements, some gene clusters were conserved and displayed lineage-specific characteristics (Fig. 6C). These synteny patterns indicate that reciprocal translocations have played a key role in shaping the evolution of *Trachurus* genomes.

Furthermore, we found that the *T. murphyi* genome shares a similar pattern of orthologous gene sets with the other *Trachurus* species, containing 18,594 common gene families (Fig. 6D), while 817 orthogroups are unique to *T. murphyi* (Fig. 6D). GO terms distribution among those unique *T. murphyi* orthogroups (3,715 genes) using WEGO2.0^65^ reveal genes involved in different biological processes, including localization, response to stimulus, metabolic processes, biological regulation, and signaling (Fig. 6E). In summary, our comparative genomic analyses highlight minimal genomic divergence and strong conservation among *Trachurus* species.

## Data Records

All genome data have been deposited in the NCBI SRA database under BioProject accession XXXXX, including PacBio HiFi reads (XXXXX to XXXXX) and Omni-C reads (XXXXX). The mitochondrial (XXXX) and nuclear genome assembly have been submitted to GenBank under the Whole Genome Shotgun project accession GCA_XXXXXX. Additionally, genome annotation files, including repeat annotation, gene annotation, CDS, and protein data, have been archived in FigShare and are available at the following link: XXXXXXX.

## Technical Validation

To evaluate the quality of our genome assembly and gene annotation, we employed several complementary approaches. First, we assessed potential contamination in the genome assembly using BlopToolKit^30^ (v4.4.3) to analyze GC content and sequencing depth. Contigs identified as contaminants were removed from the assembly (Table 1). After decontamination, we observed that nearly all GC content points clustered around 40%, indicating the absence of exogenous contamination from bacteria or other species (Fig. 1B). Second, we mapped the Omni-C reads to the assembled genome using BWA (https://github.com/lh3/bwa) to generate a Hi-C contact map. The resulting heatmap displayed high consistency across all chromosomes, providing strong evidence for the accurate sequencing, ordering, and orientation of contigs in the Chilean jack mackerel genome assembly (Fig. 1C). Third, we predicted centromeric and telomeric sequences in the *T. murphyi* genome and found that at least 12 chromosomes contained one of these characteristic chromosomal features (Fig. 2A, Table 2, Table 3), suggesting that our genome assembly qualifies as a near telomere-to-telomere assembly.

Fourth, we evaluated assembly completeness using BUSCO^34^ (v5.3.2). The analysis, based on the actinopterygii_odb10 database, revealed a completeness score of 99.0%, comprising 97.9% complete single-copy, 1.1% duplicated, 0.3% fragmented, and 0.7% missing genes (Fig. 1D). A similar result was obtained using the metazoan_odb10 database, yielding a completeness score of 98.8% distributed in 96.4% complete single-copy, 2.4% duplicated, 0.5% fragmented, and 0.7% missing genes (Fig. 1D). These results underscore the high completeness and accuracy of the Chilean jack mackerel genome assembly. Fifth, we assessed per-base consensus accuracy using Merqury^66^ (v1.3). The assembly shows very high consensus accuracy (QV = 52.8), corresponding to a base-level accuracy of 99.0005%. The low number of missing k-mers (90K out of 818 Mbp) indicates a nearly complete and high accurate assembly, with few sequence errors or gaps, indicative of a nearly error-free assembly. Sixth, we aligned PacBio HiFi reads to the assembled genome using minimap2^61^ (v2.17), resulting in a mapping rate of 99.97%, further supporting the high consensus accuracy of the assembly. Seventh, we evaluated assembly contiguity using QUAST^67^ (v5.3.0), obtaining contig N50, scaffold N50, and additional key metrics (Table 5), all demonstrating the high contiguity of the genome assembly.

**Table 5.** QUAST evaluation of the *T. murphyi* genome assembly quality.

| Assembly | Metric |
| --- | --- |
| # contigs (>= 0 bp) | 523 |
| # contigs (>= 1000 bp) | 523 |
| # contigs (>= 5000 bp) | 523 |
| # contigs (>= 10000 bp) | 523 |
| # contigs (>= 25000 bp) | 523 |
| # contigs (>= 50000 bp) | 523 |
| Total length (>= 0 bp) | 818,310,266 |
| Total length (>= 1000 bp) | 818,310,266 |
| Total length (>= 5000 bp) | 818,310,266 |
| Total length (>= 10000 bp) | 818,303,047 |
| Total length (>= 25000 bp) | 814,740,251 |
| Total length (>= 50000 bp) | 807,049,109 |
| # contigs | 523 |
| Largest contig | 43,616,551 |
| Total length | 818,310,226 |
| GC (%) | 42.31 |
| N50 | 34,335,234 |
| N90 | 25,125,839 |
| L50 | 11 |
| L90 | 22 |

Eighth, we assessed the completeness of the predicted protein-coding sequences with BUSCO^34^ (v5.3.2), achieving a BUSCO completeness score of 91.6% and 94.5% for CDS and 91.7% and 94.8% for proteins against Actinopterygii and Metazoa databases, respectively (Fig. 7). Finally, a total of 43,427 (90.98%) protein-coding genes were successfully annotated in at least one database (Fig. 4D), and 9,080 (19.02%) genes were supported by all eight databases used (Fig. 4D), indicating high-quality, accurate, and comprehensive annotation of the protein-coding genes in the *T. murphyi* genome. Taken together, all these lines of evidence confirm the accuracy, reliability, and high-quality of the Chilean jack mackerel genome assembly and annotation.

**Figure 7.**
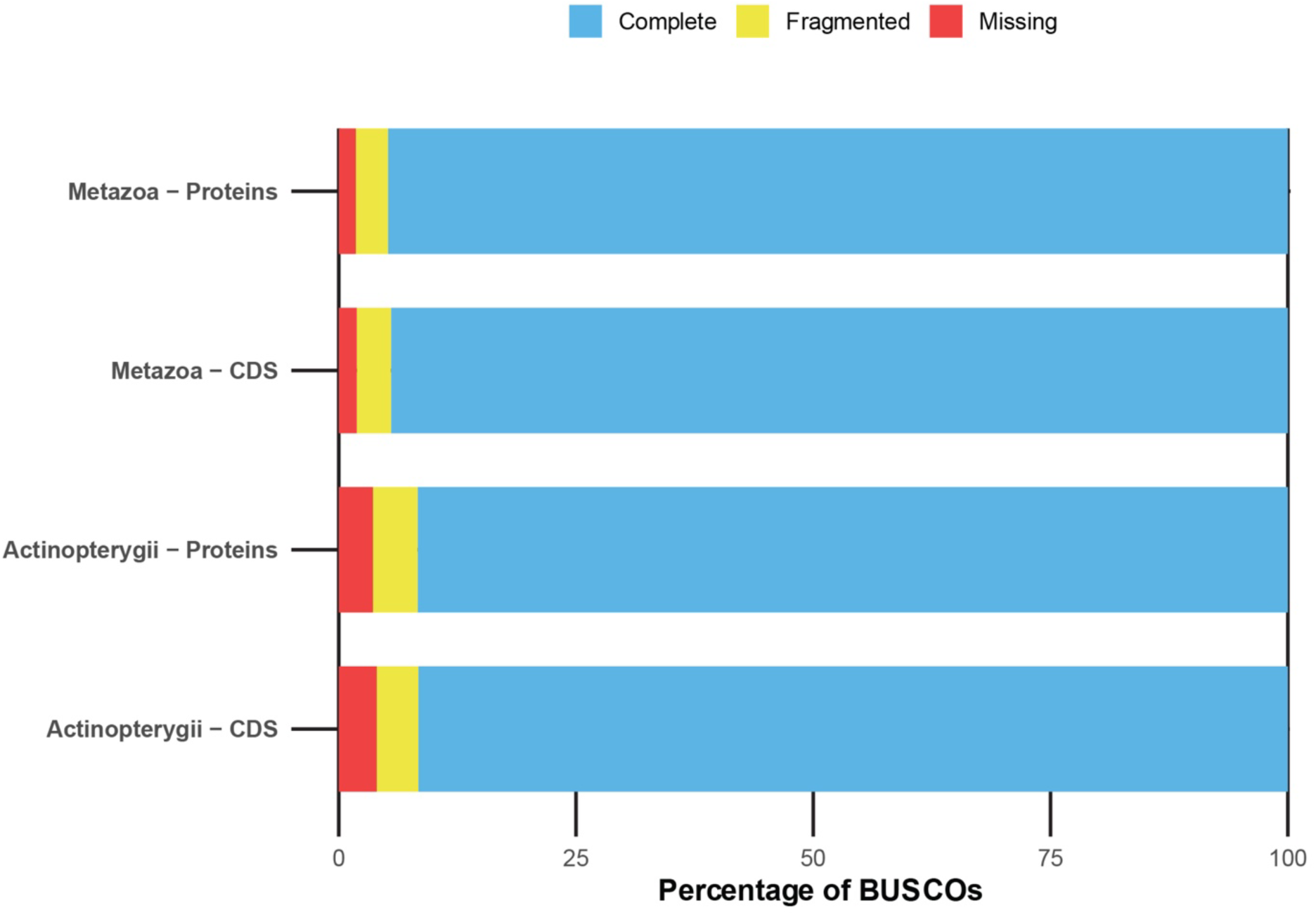
BUSCO completeness analysis of the predicted protein-coding genes in the *T. murphyi* genome assembly. The completeness of coding sequences (CDS) and protein predictions was evaluated using the BUSCO Actinopterygii and Metazoa databases.

## Code Availability

No custom codes or script were used in this study. All bioinformatics tools and pipelines were executed following the instructions and manuals provided by the respective software developers. The specific software versions and corresponding parameters, where applicable, are detailed in the Methods subsection. Unless stated otherwise, default parameters were used for all analyses.

## Acknowledgements

This work was supported by funding provided by the Fondo de Investigación Pesquera y Acuicultura of Chile, FIPA N°2021-28 and FIPA N°2023-18 to SFF, Agencia Nacional de Investigación (ANID) grants: Fondecyt Regular 1220708 and Fondequip EQM200056 to FA. CC-A received partial support from ANID Fondecyt Regular grant 1251604. We would like to thank the scientific observers from the Instituto de Investigación Pesquera (Inpesca) for collecting the Chilean jack mackerel samples.

## Author Contributions

S.F-F., C.C-A, and F.A. conceived and designed the study. S.F-F. and V.H-Y collected the Chilean jack mackerel samples and conducted laboratory procedures. C.Q. and F.A. conducted the bioinformatic analyses. F.A. drafted the initial version of the manuscript. All authors reviewed, revised, and approved the final manuscript.All authors reviewed, revised, and approved the final manuscript.

## References

1 Gushiken, S. Phylogenetic relationships of the perciform genera of the family Carangidae. Jpn J Ichthyol 34, 443–461 (1988).

2 Cárdenas, L. et al. Origin, diversification, and historical biogeography of the genus *Trachurus* (Perciformes: Carangidae). Mol Phylogenet Evol 35, 496–507 (2005).

3 Romero, J. et al. Diet and trophic position of two mackerel species in the archipielago of Madeira, Portugal. J Fish Biol 99, 831–843 (2021).

4 Gerlotto, F., Gutiérrez, M. & Bertrand, A. Insight on population structure of the Chilean jack mackerel (*Trachurus murphyi*). Aquat Living Resour 25, 341–355 (2012).

5 Serra, R. Important life history aspects of the Chilean jack mackerel, Trachurus symmetricus murphyi. Invest Pesq 36, 67–83 (1991).

6 Bertrand, A. et al. Diel vertical behaviour, predator–prey relationships, and occupation of space by jack mackerel (*Trachurus murphyi*) off Chile. ICES J Mar Sci 61, 1105–1112 (2004).

7 Bertrand, A., Barbieri, M. A., Gerlotto, F., Leiva, F. & Córdova, J. Determinism and plasticity of fish schooling behaviour as exemplified by the South Pacific jack mackerel *Trachurus murphyi*. Mar Ecol Prog Ser 311, 145–156 (2006).

8 Alegre, A. et al. Diet diversity of jack and chub mackerels and ecosystem changes in the northern Humboldt Current system: A long-term study. Prog Oceanogr 137, 299–313 (2015).

9 Bastías, J. M., Balladares, P., Acuña, S., Quevedo, R. & Muñoz, O. Determining the effect of different cooking methods on the nutritional composition of salmon (*Salmo salar*) and chilean jack mackerel (*Trachurus murphyi*) fillets. PLoS ONE 12, e0180993 (2017).

10 Lavallée, D. et al. Quebrada de los burros. Los primeros pescadores del litoral pacífico en el extremo sur peruano. Chungará (Arica) 43, 333–351 (2011).

11 Rebolledo, S., Béarez, P., Salazar, D. & Fuentes, F. Maritime fishing during the Middle Holocene in the hyperarid coast of the Atacama Desert. Quat Int 391, 3–11 (2016).

12 Arcos, D. F., Cubillos, L. A. & Núñez, S. P. The jack mackerel fishery and El Niño 1997–98 effects off Chile. Prog Oceanogr 49, 597–617 (2001).

13 Li, G., Zou, X., Chen, X., Zhou, Y. & Zhang, M. Standardization of CPUE for Chilean jack mackerel (*Trachurus murphyi*) from Chinese trawl fleets in the high seas of the Southeast Pacific Ocean. J Ocean Univ China 12, 441–451 (2013).

14 Zhu, G. et al. Does life history connectivity explain distributions of Chilean jack mackerel *Trachurus murphyi* caught in international waters prior to decline of the southeastern Pacific fishery? Fisheries Res 151, 20–25 (2014).

15 Lima, M., Canales, T. M., Wiff, R. & Montero, J. The interaction between stock dynamics, fishing and climate caused the collapse of the Jack mackerel stock at Humboldt current ecosystem. Front Mar Sci 7, 123 (2020).

16 Ashford, J., Serra, R., Saavedra, J. C. & Letelier, J. Otolith chemistry indicates large-scale connectivity in Chilean jack mackerel (*Trachurus murphyi*), a highly mobile species in the Southern Pacific Ocean. Fisheries Res 107, 291–299 (2011).

17 Cerna, F. et al. Bomb radiocarbon, otolith daily increments and length modes validate age interpretations of Chilean jack mackerel (*Trachurus murphyi*). Front Mar Sci 9, 906583 (2022).

18 Ferrada-Fuentes, S., Galleguillos, R., Herrera-Yáñez, V. & Canales-Aguirre, C. B. Population genetics of Chilean jack mackerel, *Trachurus murphyi* Nichols, 1920, (Pisces, Carangidae), in waters of the South Pacific Ocean. Fishes 8, 162 (2023).

19 Oliva, M. E. Metazoan parasites of the jack mackerel *Trachurus murphyi* (Teleostei, Carangidae) in a latitudinal gradient from South America (Chile and Peru). Parasite 6, 223–230 (1999).

20 Vásquez, S., Correa-Ramírez, M., Parada, C. & Sepúlveda, A. The influence of oceanographic processes on jack mackerel (*Trachurus murphyi*) larval distribution and population structure in the southeastern Pacific Ocean. ICES J Mar Sci 70, 1097–1107 (2013).

21 Canales-Aguirre, C. B., Ferrada, S. & Galleguillos, R. Isolation and characterization of microsatellite loci for the jack mackerel (Trachurus murphyi Nicols, 1920). Conserv Genet 11, 1235–1237 (2010).

22 Cárdenas, L. et al. Genetic population structure in the Chilean jack mackerel, Trachurus murphyi (Nichols) across the South-eastern Pacific Ocean. Fisheries Res 100, 109–115 (2009).

23 Galleguillos, R., Canales-Aguirre, C. B. & Ferrada, S. Genetic variability in jack mackerel *Trachurus murphyi* Nichols: new SSRs loci and application. Gayana (Concepción) 76, 58–62 (2012).

24 Canales-Aguirre, C. B., Ferrada-Fuentes, S. & Galleguillos, R. Temporal and spatial population genetic variation in Chilean jack mackerel (*Trachurus murphyi*). Biology 14, 510 (2025).

25 Canales-Aguirre, C. B. et al. Heterogametic females reveal a ZW sex determination system and a putative sex chromosome for the Chilean jack mackerel, *Trachurus murpyhi*. Mol Ecol Resour, e70034 (2025).

26 Asorey, C. M., Larraín, M. A. & Araneda, C. The complete mitochondrial genome of Chilean jack mackerel, *Trachurus murphyi* Nichols, 1920 (Teleostei, Carangidae). Mitochondrial DNA 9, 1455–1459 (2024).

27 Genner, M. & Collins, R. The genome sequence of the Atlantic horse mackerel, *Trachurus trachurus* (Linnaeus 1758). Wellcome Open Res 7, 118 (2022).

28 Hirao, A. S. et al. Genome-wide SNP analysis coupled with geographic and reproductive-phenological information reveals panmixia in a classical marine species, the Japanese jack mackerel (*Trachurus japonicus*). Fisheries Res 279, 107146 (2024).

29 Cheng, H., Concepcion, G. T., Feng, X., Zhang, H. & Li, H. Haplotype-resolved de novo assembly using phased assembly graphs with hifiasm. Nat Methods 18, 170–175 (2021).

30 Challis, R., Richards, E., Rajan, J., Cochrane, G. & Blaxter, M. BlobToolKit – Interactive quality assessment of genome assemblies. G3 Genes Genomes Genet 10, 1361–1374 (2020).

31 Guan, D. et al. Identifying and removing haplotypic duplication in primary genome assemblies. Bioinformatics 36, 2896–2898 (2020).

32 Putnam, N. H. et al. Chromosome-scale shotgun assembly using an in vitro method for long-range linkage. Genome Res 26, 342–350 (2016).

33 Durand, N. C. et al. Juicebox provides a visualization system for Hi-C contact maps with unlimited zoom. Cell Syst 3, 99–101 (2016).

34 Manni, M., Berkeley, M. R., Seppey, M., Simão, F. A. & Zdobnov, E. M. BUSCO update: Novel and streamlined workflows along with broader and deeper phylogenetic coverage for scoring of eukaryotic, prokaryotic, and viral genomes. Mol Biol Evol 38, 4647–4654 (2021).

35 Gomes, N. M. V., Shay, J. W. & Wright, W. E. Telomere biology in Metazoa. FEBS Lett 584, 3741–3751 (2010).

36 Lin, Y. et al. quarTeT: a telomere-to-telomere toolkit for gap-free genome assembly and centromeric repeat identification. Hortic Res 10, uhad127 (2023).

37 Miga, K. H. Completing the human genome: the progress and challenge of satellite DNA assembly. Chromosome Res 23, 421–426 (2015).

38 Rocchi, M., Archidiacomo, N., Schempp, W., Capozzi, O. & Stanyon, R. Centromere repositioning in mammals. Heredity 108, 59–67 (2012).

39 Pei, T. et al. Gap-free genome assembly and CYP450 gene family analysis reveal the biosynthesis of anthocyanins in *Scutellaria baicalensis*. Hortic Res 10, uhad235 (2023).

40 Li, M. et al. Telomere-to-telomere genome assembly of sorghum. Sci Data 11, 835 (2024).

41 Flynn, J. M. et al. RepeatModeler2 for automated genomic discovery oftransposable element families. Proc Natl Acad Sci USA 117, 9451–9457 (2020).

42 Haas, B. TransposonPSI: an application of PSI-Blast to mine (retro-) transposon ORF homologies. https://transposonpsi.sourceforge.net/ (2007).

43 Ellinghaus, D., Kurtz, S. & Willhoeft, U. LTRharvest, an efficient and flexible software for de novo detection of LTR retrotransposons. BMC Bioinformatics 9, 18 (2008).

44 Steinbiss, S., Willhoeft, U., Gremme, G. & Kurtz, S. Fine-grained annotation and classification of de novo predicted LTR retrotransposons. Nucleic Acids Res 37, 7002–7013 (2009).

45 Gremme, G., Steinbiss, S. & Kurtz, S. GenomeTools: A aomprehensive software library for efficient processing of structured genome annotations. IEEE/ACM Trans Comput Biol Bioinform 10, 645–656 (2013).

46 Crescente, J. M., Zavallo, D., Helguera, M. & Vanzetti, L. S. MITE Tracker: an accurate approach to identify miniature inverted-repeat transposable elements in large genomes. BMC Bioinformatics 19, 348 (2018).

47 Xiong, W., He, L., Lai, J., Dooner, H. K. & Du, C. HelitronScanner uncovers a large overlooked cache ofHelitron transposons in many plant genomes. Proc Natl Acad Sci USA 111, 10263–10268 (2014).

48 Edgar, R. C. Search and clustering orders of magnitude faster than BLAST. Bioinformatics 26, 2460–2461 (2010).

49 Camacho, C. et al. BLAST+: architecture and applications. BMC Bioinformatics 10, 421, doi:10.1186/1471-2105-10-421 (2009).

50 Tarailo-Graovac, M. & Chen, N. Using RepeatMasker to identify repetitive elements in genomic sequences. Curr Protoc Bioinformatics 4, 4.10.11-14.10.14 (2009).

51 Gabriel, L. et al. BRAKER3: Fully automated genome annotation using RNA-seq and protein evidence with GeneMark-ETP, AUGUSTUS, and TSEBRA. Genome Res 34, 769–777 (2024).

52 Bruna, T., Lomsadze, A. & Borodovsky, M. GeneMark-ETP significantly improves the accuracy of automatic annotation of large eukaryotic genomes. Genome Res 34, 757–768 (2024).

53 Stanke, M. & Morgenstern, B. AUGUSTUS: a web server for gene prediction in eukaryotes that allows user-defined constraints. Nucleic Acids Res 33, W465–W467 (2005).

54 Gabriel, L., Hoff, K. J., Bruna, T., Borodovsky, M. & Stanke, M. TSEBRA: transcript selector for BRAKER. BMC Bioinformatics 22, 566 (2021).

55 Buchfink, B., Reuter, K. & Drost, H.-G. Sensitive protein alignments at tree-of-life scale using DIAMOND. Nat Methods 18, 366–368 (2021).

56 Johnson, L. S., Eddy, S. R. & Portugaly, E. Hidden Markov model speed heuristic and iterative HMM search procedure. BMC Bioinformatics 11, 431 (2010).

57 Jones, P. et al. InterProScan 5: genome-scale protein function classification. Bioinformatics 30, 1236–1240 (2014).

58 Aramaki, T. et al. KofamKOALA: KEGG ortholog assignment based on profile HMM and adaptive score threshold. Bioinformatics 36, 2251–2252 (2020).

59 Uliano-Silva, M. et al. MitoHiFi: a python pipeline for mitochondrial genome assembly from PacBio high fidelity reads. BMC Bioinformatics 24, 288 (2023).

60 Donath, A. et al. Improved annotation of protein-coding genes boundaries in metazoan mitochondrial genomes. Nucleic Acids Res 47, 10543–10552 (2019).

61 Li, H. Minimap2: pairwise alignment for nucleotide sequences. Bioinformatics 34, 3094–3100 (2018).

62 Grant, J. R. et al. Proksee: in-depth characterization and visualization of bacterial genomes. Nucleic Acids Res 51, W484–W492 (2023).

63 Cabanettes, F. & Kloop, C. D-GENIES: dot plot large genomes in an interactive, efficient and simple way. PeerJ 6, e4958 (2018).

64 Emms, D. M. & Kelly, S. OrthoFinder: phylogenetic orthology inference for comparative genomics. Genome Biol 20, 238 (2019).

65 Ye, J. et al. WEGO 2.0: A web tool for analyzing and plotting GO annotations, 2018 update. Nucleic Acids Res 46, W71–W75 (2018).

66 Rhie, A., Walenz, B. P., Koren, S. & Phillippy, A. M. Merqury: reference-free quality, completeness, and phasing assessment for genome assemblies. Genome Biol 21, 245 (2020).

67 Gurevich, A., Saveliev, V., Vyahhi, N. & Tesler, G. QUAST: quality assessment tool for genome assemblies. Bioinformatics 29, 1072–1075 (2013).

